# Elevated glucocorticoid alters the developmental dynamics of hypothalamic neurogenesis

**DOI:** 10.1101/2023.01.27.525966

**Authors:** Helen Eachus, Min-Kyeung Choi, Anna Tochwin, Johanna Kaspareit, May Ho, Soojin Ryu

## Abstract

Exposure to excess glucocorticoid (GC) during early development is implicated in adult dysfunctions. Reduced adult hippocampal neurogenesis is a well-known consequence of exposure to early life stress or elevated GC, however the effects on neurogenesis during development and effects on other brain regions are not well understood. Using an optogenetic zebrafish model, here we analysed the effects of GC exposure on neurogenesis during development in the whole brain. We identify that the hypothalamus is a highly GC-sensitive region where elevated GC causes precocious development. This is followed by failed maturation and early decline accompanied by impaired feeding, growth, and longevity. In GC-exposed animals, the developmental trajectory of hypothalamic progenitor cells is strikingly altered, potentially mediated by direct regulation of transcription factors such as *rx3* by GC. Our data provide cellular and molecular level insight into GC-induced alteration of the hypothalamic development trajectory, a process crucial for health across the life-course.

## Introduction

Glucocorticoids (GC) are the key effectors of the stress response and have pleiotropic effects on the body, acting to restore homeostasis and thus allow an animal to respond adaptively to threat^1^. Since the developing brain is plastic, GC exposure during early life has the potential to alter the developmental trajectory of the brain. Indeed, alteration of the development trajectory may be an adaptive mechanism employed by animals exposed to stress during early life. For example, antenatal GC treatment has well documented effects on lung maturation, as well as other organs, in preterm babies^2^. However, such treatment has also been associated with negative outcomes in later life, such as increased incidence of mental health issues during childhood^3^, impacted development of fronto-parietal brain functions during adolescence ^4^, and cortical thinning in children with associated affective disorders^5^. Early life GC exposure leading to adverse consequences in later life is an example of the concept known as early life programming of adult disease^6–9^.

One of the well-documented effects of GC exposure during early life or early life stress (ELS) is reduced adult neurogenesis in the hippocampus, including reduced cell proliferation^10–14^. Indeed adrenalectomy is sufficient to prevent reduced proliferation in chronically stressed mice, suggesting that GC is the main driving force behind this phenotype^15^. In contrast, only a few studies have looked at the effects of ELS or GC exposure on neurogenesis during development, but there are some indications that its effects might be developmentally dynamic. Specifically, it is possible that GC-induced reduction of cell proliferation is a delayed consequence of a GC-induced change to the development trajectory that began in early life. Indeed, rats exposed to ELS exhibited enhanced hippocampal cell proliferation and improved stress-associated behavioural performance as young adults, whilst these effects were reversed by middle age^10^. A potential mechanistic explanation for the ELS-induced reduction in adult hippocampal neurogenesis is that enhanced cell proliferation at an earlier time-point depletes the stem cell pool over time^16^.

Despite the well-documented effects of GC exposure on hippocampal neurogenesis, the effects of GC exposure on other brain regions are unclear. Recently, a first study of the effects of ELS on hypothalamic neurogenesis revealed that cell proliferation and numbers of hypothalamic stem cells known as tanycytes were reduced in the adult mouse^17^. Tanycytes are unique radial glial-like cells that line the walls of the 3^rd^ ventricle, have NSPC (neural stem/progenitor cell) properties that persist into adulthood, send projections into neighbouring hypothalamic nuclei and are able to sense glucose levels in the CSF^18^. Whilst most previous studies have focused on effects of stress and GC exposure on neurogenesis in the hippocampus, the aforementioned study supports that the hippocampus is not uniquely affected by stress/GC. However, whether and how GC affects hypothalamic neurogenesis during development is completely unknown.

Here we used an optogenetic zebrafish model in which endogenous cortisol level is elevated to test the hypothesis that GC exposure alters the developmental trajectory of the brain. The model utilises optogenetic manipulation of steroidogenic interrenal cells to increase the endogenous GC level during larval and juvenile stages^19^. Whilst the mammalian adrenal gland, counterpart of the fish interrenal, is known to produce GCs, mineralocorticoids (MCs), and androgens^20^; MCs and androgens are not produced by the larval zebrafish interrenal gland^21–24^. Therefore, the phenotypes observed in our model are primarily caused by GC and its downstream effectors, rather than MC or androgen over-exposure.

We found that GC exposure affected cell proliferation in a region-specific manner and that these effects were dynamic across the life-course. We found that the hypothalamus is a highly GC-sensitive brain region. In early life, we observed precocious development of the hypothalamus and hypothalamus-associated feeding behaviour in GC-exposed animals. This was followed by a rapid decline, indicated by failed hypothalamic maturation, including reduced numbers of *agrp*- and *hcrt*-expressing neurons known to regulate feeding^25–27^, and suppressed feeding and growth. Our data uncover a tissue-specific time-dependent plasticity of the hypothalamus in response to GC and provide cellular and molecular insights for GC-induced alteration of the hypothalamic development trajectory.

## Results

### Neurogenesis is altered in an optogenetic zebrafish model of elevated GC

Our lab previously generated an optogenetic zebrafish model, *Tg(star:bPAC-2A-tdTomato)^uex^*^300^, in which the endogenous cortisol level can be increased specifically in response to blue light exposure using an optogenetic protein *beggiatoa* Photoactivated Adenylate Cylase (bPAC) (Fig.1a)^19, 28, 29^. In order to determine the effects of elevated GC on the brain across development, we raised transgenic fish under standard aquarium lighting conditions, where the blue light component of the white light is sufficient to activate bPAC, which led to significantly elevated cortisol levels in *Tg(star:bPAC-2A-tdTomato)^uex^*^300^ compared with wild types from hatching through until late juvenile/early adult stage (Fig.1b)^19^. *Tg(star:bPAC-2A-tdTomato)^uex^*^300^ raised under standard light conditions will be referred to throughout as star:bPAC+ or bPAC+.

**Figure 1.**
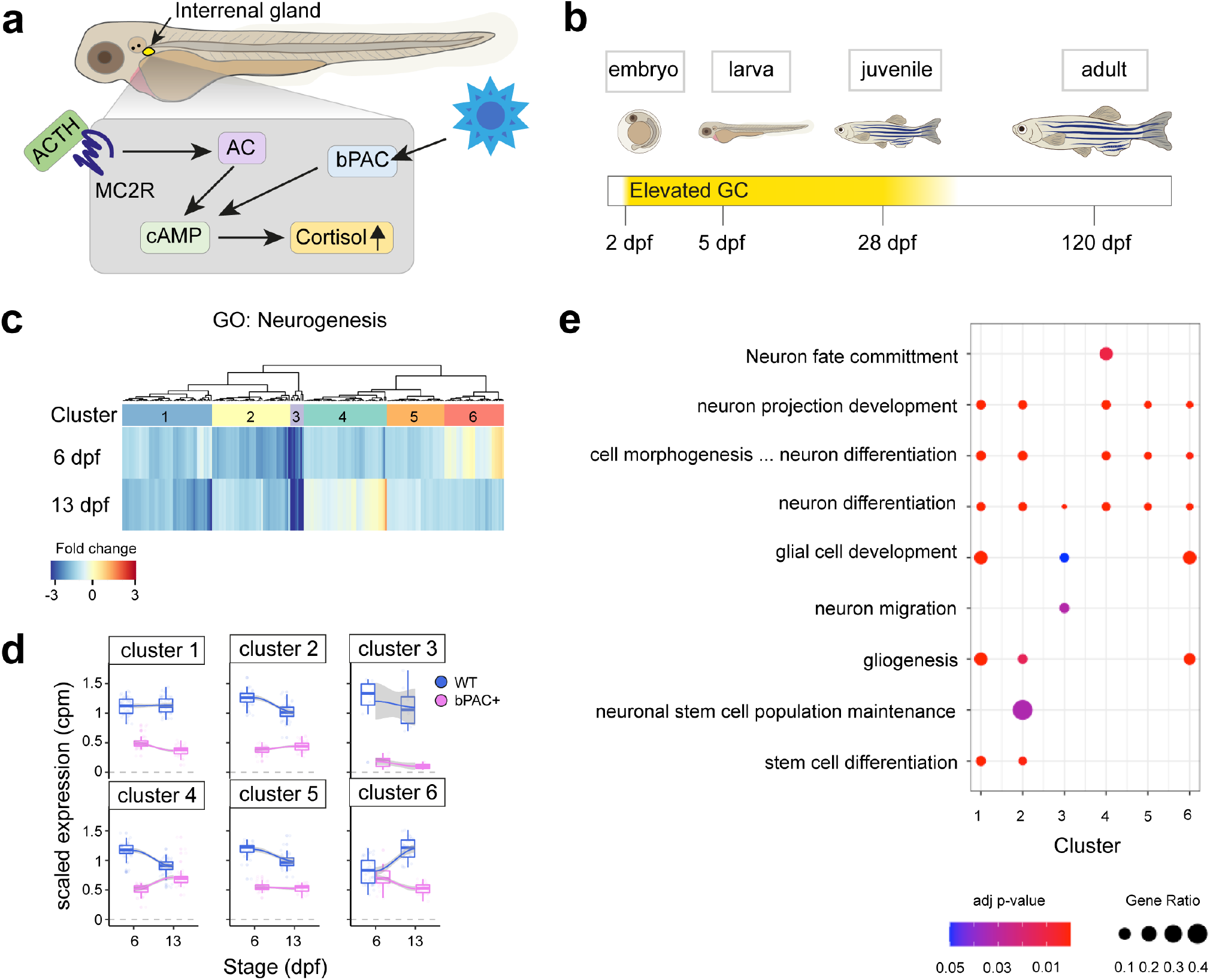
Neurogenesis is altered in an optogenetic zebrafish model of elevated GC. **a**. In star:bPAC+ larvae, blue light activates cAMP signalling in the interrenal gland, which leads to release of cortisol. Figure adapted from Eachus et al, 2021^81^. **b.** Wild-type and star:bPAC+ larvae were raised under a standard 12:12 light/dark cycle under white light. In these conditions, basal cortisol levels are elevated in star:bPAC+ larvae from 2 dpf until approximately 2 months of age. **c.** Heatmap showing fold change for 225 genes associated with GO:0022008 (neurogenesis) in star:bPAC+ whole-brain samples that were downregulated at either 6 dpf or 13 dpf (or both), relative to wild type, from an RNA-sequencing analysis. **d.** Scaled expression (cpm) of neurogenesis-associated genes in star:bPAC+ and wild-type whole-brain samples at 6 dpf and 13 dpf, according to the 6 clusters shown in c. Boxes show the median and quartiles, whiskers are inter-quartile range and gray shading shows the smoothed conditional means. **e**. Significant slimmed neurogenesis child GO terms at p<0.05, identified using g:Profiler, are shown for each cluster. Size of the dot indicates the ratio of differentially expressed genes from each GO term, whilst the colour shows the adjusted p-value.

To investigate the effects of elevated GC on brain development we performed bulk RNA-sequencing (RNA-seq) of whole brain samples from star:bPAC+ and wild-type fish at different developmental stages^19^. Gene Ontology (GO) analysis of differentially expressed genes (DEGs) that were down-regulated in star:bPAC+ brains compared with wild types were enriched for a number of biological processes, including Neurogenesis at all developmental stages (GO:0022008, adjusted p-value 2.47E-20 at 6 days post fertilization (dpf), 5.45E-08 at 13 dpf, 1.11E-17 at 120 dpf). Up-regulated genes were not enriched for GO term neurogenesis. To investigate the effects of GC exposure on neurogenesis during development, we focused downstream analysis on the early life time-points: 5/6 dpf and 13 dpf. 225 genes associated with neurogenesis were down-regulated in star:bPAC+ brains at either 6 dpf or 13 dpf (Fig.1c). To confirm our results and identify specific features of neurogenesis that are altered by elevated GC, we performed qPCR on independent samples of whole brains from 5 dpf star:bPAC+ and wild types for genes associated with neurogenesis processes or distinct cell types. Of the 14 categories of genes analysed, only one category was associated with consistent differential expression in star:bPAC+. Expression of *pcna*, *mki67* and *mcm5*, all associated with cell proliferation, showed increased expression in star:bPAC+ compared with wild types (Supplementary Fig.1). These data support that neurogenesis, and especially cell proliferation, is altered by elevated GC.

To identify whether the effect of GC on neurogenesis is developmentally dynamic, we performed unsupervised clustering analysis of neurogenesis DEGs at 6 dpf and 13 dpf and identified 6 different clusters. Cluster 2 and cluster 5 genes showed a relatively stronger down-regulation at 6 dpf compared to 13 dpf, meanwhile cluster 1 genes showed a relatively stronger downregulation at 13 dpf (Fig.1d). Cluster 3 genes were strongly downregulated at both time-points (Fig.1d). Interestingly, cluster 4 and cluster 6 genes showed a timepoint-specific down-regulation. Cluster 4 genes were downregulated in star:bPAC+ at 6 dpf but were not differentially expressed at 13 dpf (Fig.1d). Meanwhile cluster 6 genes were down-regulated in star:bPAC+ at 13 dpf but not at 6 dpf (Fig.1d). These results suggest that some effects of elevated GC on neurogenesis are temporarily dynamic during early life. GO analysis of genes within the clusters indicated that all clusters contained genes associated with neuron differentiation (Fig.1e). Gliogenesis and glial cell development were enriched in multiple clusters. There were also some terms which were restricted to single or small numbers of clusters. For example, cluster 4 genes were associated with neuron fate commitment, cluster 3 with neuron migration, and cluster 2 with neuronal stem cell population maintenance (Fig.1e). These results suggest that specific aspects of neurogenesis exhibit temporally dynamic differential expression during early life as a result of GC exposure.

### Precocious hypothalamic development following elevated GC

In order to determine whether the effects of elevated GC on cell proliferation were brain-wide or region specific we performed immunohistochemistry (IHC) for mitosis marker phospho-histoneH3 (pH3) on brains of 5 dpf star:bPAC+ and wild-type larvae (Fig.2a). Cell counting of proliferating pH3+ cells across brain regions indicated a trend towards an increase in number of pH3+ cells in the valvula cerebelli (ce-v) of the hindbrain, however, the only significant increase was restricted to the hypothalamus (Fig.2b). We next sought to determine which cell types were associated with the increased proliferative capacity in the hypothalamus of 5 dpf star:bPAC+. To this end, we performed Fluorescent *in situ* hybridization (FISH) for *rx3* (*retinal homeobox gene 3),* combined with IHC for Pcna and pH3. *rx3* is a transcription factor known to be expressed in hypothalamic progenitor cells in zebrafish where it plays a critical role in hypothalamic development^30^, and its mammalian orthologue *Rax* is known to be expressed in hypothalamic radial glia^31^. Pcna and pH3 are markers for proliferating cells during S-phase and M-phase respectively. Pcna+ cells were situated around the 3^rd^ ventricle of the hypothalamus, especially around the lateral recess, in both star:bPAC+ and wild-type larvae, with a small number of pH3+ cells scattered throughout this region (Fig.2c). We distinguish the Pcna+ region around the hypothalamic ventricles as the ‘proliferative zone’ and observed co-localisation of *rx3* with Pcna and pH3 within the proliferative zone (Fig.2c, white arrowheads).

**Figure 2.**
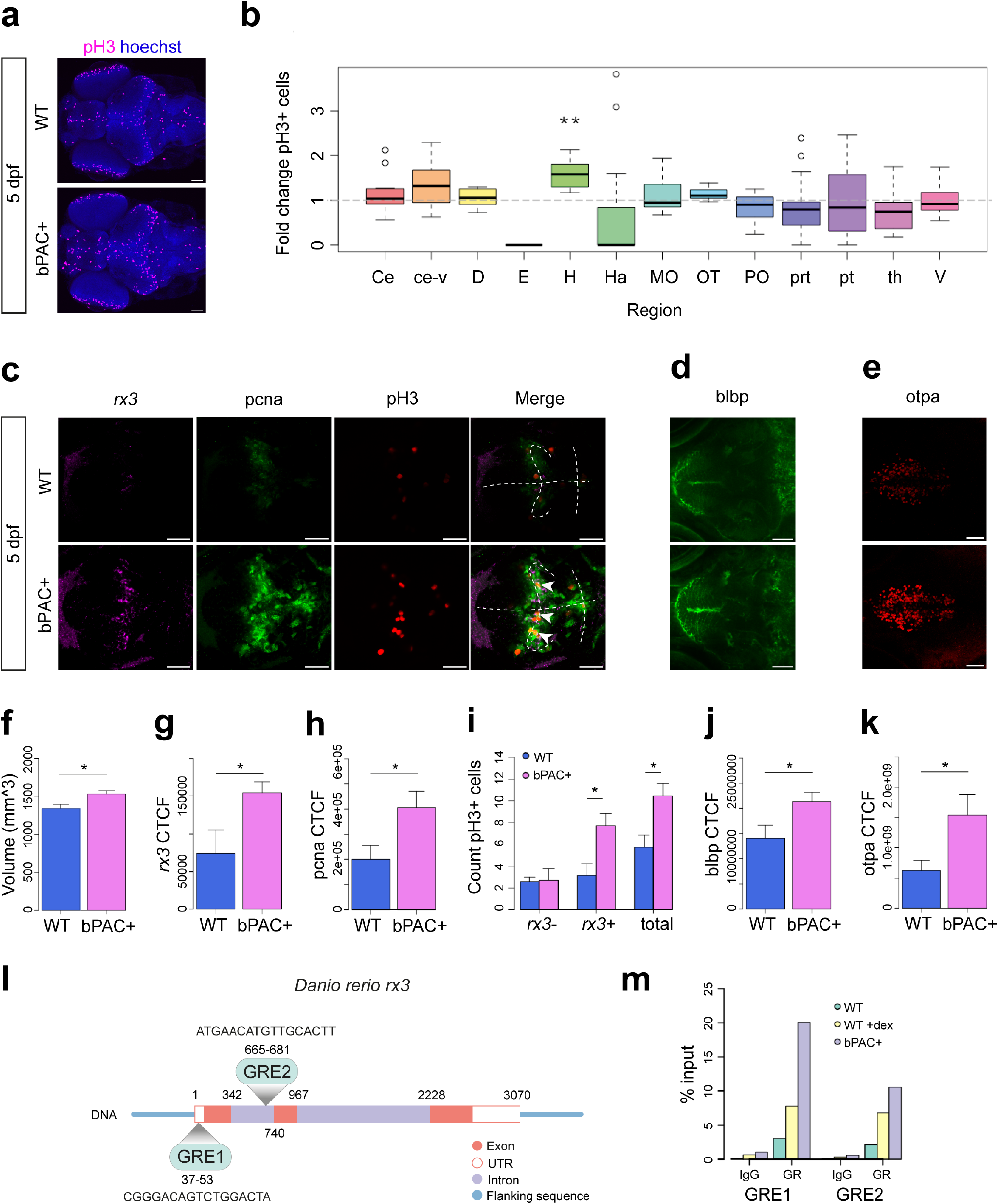
Precocious hypothalamic development following elevated GC. **a.** Whole-brain IHC for proliferation marker pH3 as seen in a maximum intensity projection in 5 dpf wild-type and star:bPAC+ larvae. Scale bar is 60 μm. **b.** Cell counting of pH3 across brain regions revealed an increase in the hypothalamus (H) of 5 dpf star:bPAC+ larvae, compared with wild type (Wilcox Exact test with Bonferroni correction (N=14, p= 0.0013195). Brain regions are labelled according to the ZFIN anatomy database. **c**. Co-expression of *rx3*, pcna and pH3 in the hypothalamus of 5 dpf wild-type and star:bPAC+ larvae, observed in a single plane image of FISH-IHC. Scale bar 40 μm. Dashed line indicates the ventricles and arrowheads indicate excess *rx3*+ pH3+ cells in star:bPAC+ hypothalamus. **d.** IHC for blbp in the arcuate nucleus of 5 dpf wild-type and star:bPAC+ larvae, observed in a single plane image. Scale bar 40 μm**. e.** IHC for otpa in the hypothalamus of 5 dpf wild-type and star:bPAC+ larvae, as observed in a maximum intensity projection. Scale bar 40 μm**. f-h**. The hypothalamus of 5 dpf star:bPAC+ larvae is larger (f, t= -2.6193, df= 11.207, p-value = 0.02353) and has increased intensity of *rx3* (g, N=7, t= -2.3011, df= 8.5909, p-value = 0.04823) and pcna (h, N=7, t =-2.4409, df= 11.766, p-value = 0.03147). **i.** Excess proliferating pH3+ cells in the hypothalamus of 5 dpf star:bPAC+ larvae are *rx3*+ radial glia (N=7, Two-way ANOVA with Tukey’s test: total pH3+ cells (p= 0.0293550), *rx3*+ pH3+ cells (p= 0.0373125)). **j-k.** Intensity of blbp (j, N=9, t= -2.2734, df= 14.903, p-value= 0.03823) and otpa (k, N=12, t= -2.44, df= 16.227, p-value= 0.02652) was increased in the hypothalamus of 5 dpf star:bPAC+ larvae. **l.** The zebrafish *rx3* gene contains two GREs, one in exon one and another in intron one. The region shown is ENSDARG00000052893, GRCz11:21:10755554:10759823:1 (including flanking sequence). **m.** ChIP qPCR analysis of two GREs in the *rx3* gene shows enrichment for GR compared with IgG control in 5 dpf wild-type brains. Enrichment of GR was higher in wild types exposed to dexamethasone, and higher still in star:bPAC+ brains for both GREs.

Quantitative analysis of confocal stacks revealed that the hypothalamus was significantly larger (Fig.2f), and we found a significant increase in *rx3* and pcna signal intensity, and significantly more pH3+ cells in the hypothalamus of 5 dpf star:bPAC+ compared with wild types (Fig.2g-i). To determine whether the observed increase in hypothalamic proliferation in the hypothalamus of 5 dpf star:bPAC+ larvae is a direct effect of developmental GC exposure, we treated star:bPAC+ larvae with the GR antagonist RU-486 from 2 dpf until 5 dpf. Treatment with RU-486 did not significantly affect hypothalamic volume and elicited only a trend reduction in hypothalamic Pcna, however it was sufficient to significantly reduce the number of pH3+ mitotic cells in the hypothalamus (Supplementary Fig.2), suggesting that GR signalling is a significant driver of the increased hypothalamic proliferation observed in star:bPAC+ larvae. Further, we observed that the excess hypothalamic pH3+ cells were *rx3*+, meanwhile *rx3*-/pH3+ cells were not different (Fig.2i). These results suggest that excess proliferating hypothalamic cells in star:bPAC+ are *rx3*-expressing radial glia. Interestingly, we identified two Glucocorticoid Response Elements (GREs) within the zebrafish *rx3* gene (Fig.2l). GRE1 is in the 5’ UTR, which is most likely a regulatory region. ChIP-qPCR analysis revealed enrichment of GR antibody (compared with IgG) at both GREs in larval brain samples, supporting that GR does indeed bind to regulatory regions of the *rx3* gene (Fig.2m). Further, enrichment of GR was higher in dexamethasone-treated wild-type samples, compared with wild-type controls, and higher still in star:bPAC+ samples. This experiment shows that GC exposure directly regulates GR-mediated control of *rx3* gene expression in the brain.

Whilst *rx3* appeared to be co-expressed with Pcna in proliferating radial glia, we also analysed Blbp (brain lipid binding protein, fabp7a) localisation, which is known to be expressed in predominantly quiescent radial glia in the zebrafish brain^32^. Consistent with this, we observed strong expression of Blbp in cells lining the third ventricle in the Pcna-negative rostral domains of the hypothalamus, whilst some Blbp signal was also observed in Pcna-positive cells of the lateral recess (Fig.2d). In 5 dpf star:bPAC+ larvae, we observed increased Blbp signal, especially in the arcuate region of the hypothalamus (Fig.2d,j), suggesting that increased numbers of glial cells might include both proliferative and quiescent subtypes. Finally we observed increased signal for Otpa (orthopedia homeobox a) in the hypothalamus of 5 dpf star:bPAC+ larvae (Fig.2e,k), which is predominantly expressed in early post-mitotic neuronal precursors^33^, suggesting that the excess GC-exposed hypothalamic proliferative radial glia do exit the cell cycle, leading to increased numbers of neuronal precursors. Together these data suggest that elevated GC drives precocious development of the hypothalamus in zebrafish larvae.

### Failed hypothalamic maturation following elevated GC

To test whether the effects of GC exposure on hypothalamic neurogenesis might be temporally dynamic, we investigated hypothalamic cell proliferation at 13 dpf. In the 13 dpf hypothalamus, *rx3* was expressed in a subset of Pcna+ cells around the lateral recess, and Blbp was expressed mainly along the midline 3^rd^ ventricle anterior to the proliferative zone, in a similar manner to at 5 dpf (Fig.3a). In 13 dpf star:bPAC+ larvae, we observed that the size of the hypothalamus was no longer larger, which was previously observed at 5 dpf (Fig.3b). Expression of hypothalamic *rx3* was low at this developmental stage, and showed a trend towards a reduction in star:bPAC+ compared with wild type (Fig.3a,c). In stark contrast to the 5 dpf hypothalamus, we now observed a dramatic reduction in expression of Pcna in 13 dpf star:bPAC+ hypothalamus (Fig.3a,d). Meanwhile, Blbp signal intensity was not different between 13 dpf wild-type and star:bPAC+ larvae (Fig.3a,e).

**Figure 3.**
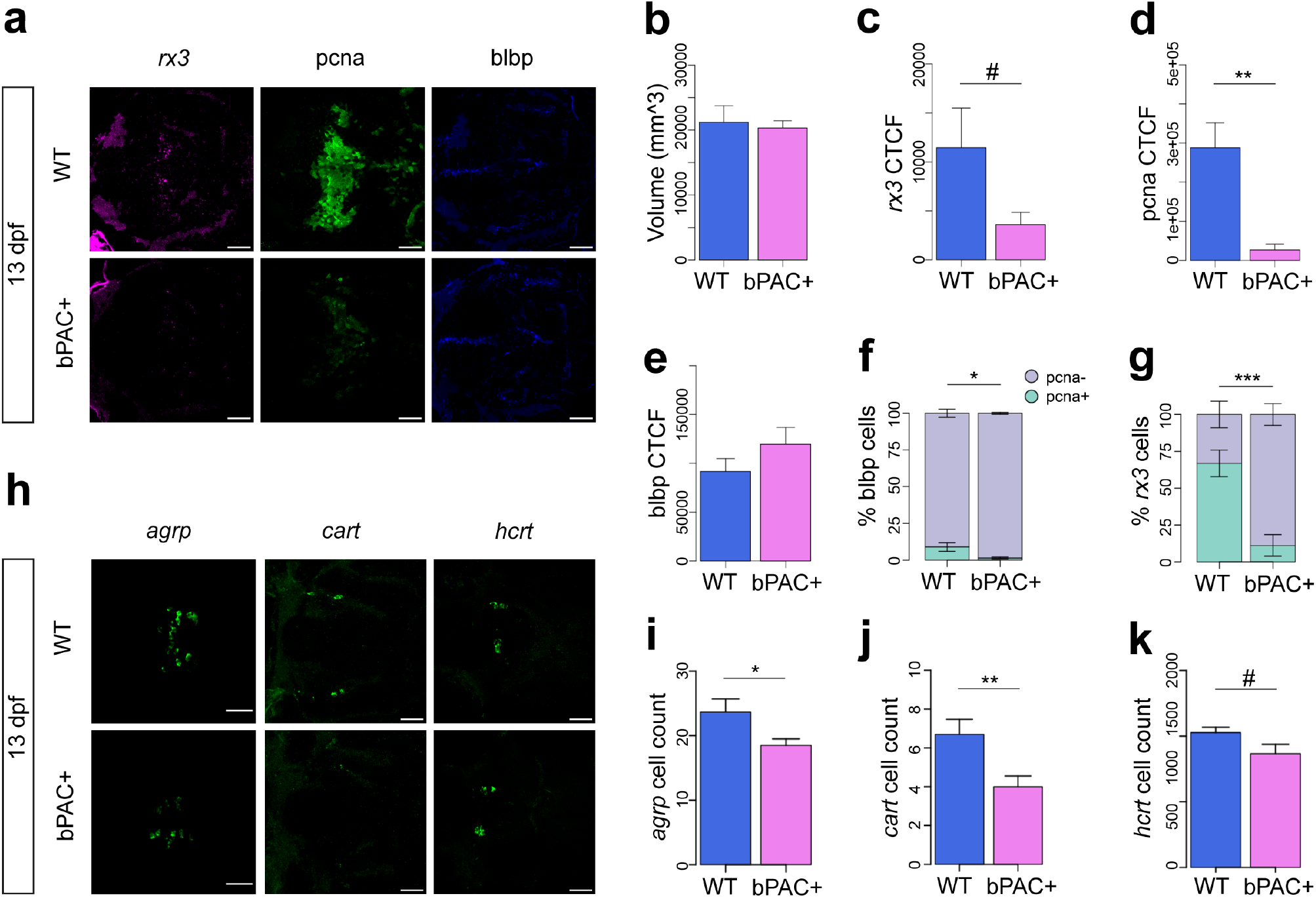
Failed hypothalamic maturation following elevated GC. **a**. Expression of *rx3*, pcna and blbp in the hypothalamus of 13 dpf WT and star:bPAC larvae, observed via FISH-IHC in single plane confocal images through the hypothalamus. Scale bar is 40 μm. **b-e.** Quantitative analysis of confocal images shown in (a), N=8. Volume of the hypothalamus is not different (b, t = 0.31771, df = 10.899, p-value = 0.7567), meanwhile, *rx3* CTCF shows a trend towards reduction (c, t = 1.8613, df = 8.3108, p-value = 0.09835), pcna CTCF is significantly reduced (d, t = 4.0207, df = 7.7651, p-value = 0.00408), and blbp CTCF is not significantly different (e, t = -1.2827, df = 13.092, p-value = 0.2219) in the hypothalamus of 13 dpf star:bPAC+ larvae compared with wild types. **f-g.** Quantitative analysis of co-expression of pcna in hypothalamic radial glia, from the images shown in (a). blbp cells (f, t = -2.5888, df = 7.5126, p-value = 0.0339) and *rx3* cells (g, t = 4.7867, df = 13.444, p-value = 0.0003238) have a significant reduction in pcna co-expression in the hypothalamus of 13 dpf star:bPAC+ larvae compared with wild types. **h.** Expression of genes involved in feeding regulation in the hypothalamus of 13 dpf star:bPAC+ and wild-type larvae as shown in maximum intensity projections of confocal stacks. Scale bar is 40 μm. **i-k.** Quantitative analysis of cell counts in the stacks shown in (h). Number of *agrp* cells is significantly lower (i, N=6-9, t = 2.2891, df = 11.255, p-value = 0.04234), number of *cart4* cells is significantly reduced (j, N=20-21, t = 2.8544, df = 34.862, p-value = 0.007212), number of *hcrt* cells shows a trend reduction (k, N=15-18, t = 1.9553, df = 26.571, p-value = 0.06115). All graphs show the mean with standard error. #, p<0.1, *, p<0.05, **, p<0.01, ***, p<0.001.

Strikingly, when we performed co-expression analysis, we observed that whilst only a small proportion of Blbp+ cells were proliferative in 13 dpf wild-type hypothalamus (∼10%), almost all Blbp+ cells were quiescent in star:bPAC+ (Fig.3f). Further, we observed that in wild types most hypothalamic *rx3+* cells (∼70%) were proliferative and expressed Pcna, however in star:bPAC+ most *rx3+* cells (∼85%) were quiescent and didn’t express Pcna (Fig.3g). These data support that proliferative radial glia are lost in the hypothalamus of 13 dpf star:bPAC+ larvae. Next, we sought to identify an explanation for the loss of proliferative radial glia in the hypothalamus. We first analysed cell death in the hypothalamus of 13 dpf larvae using a TUNEL assay. We did not observe a significant difference in the number of apoptotic cells in the hypothalamus of 13 dpf star:bPAC+ larvae compared to wild types (Supplementary Fig.3a), suggesting that loss of hypothalamic radial glia is not due to increased cell death. We reasoned that the loss of proliferation in the star:bPAC+ hypothalamus might be due to senescence of the formerly highly proliferative radial glia. In support of this hypothesis, we observed that one of the five significantly enriched KEGG pathways in our whole brain RNA-seq data set in the 13 dpf star:bPAC+ was cellular senescence, a pathway which was not enriched in the 6 dpf star:bPAC+ samples (Supplementary Fig.3b,c). In the RNA-seq analysis, 102 of 150 genes associated with the KEGG pathway cellular senescence were differentially expressed in 13 dpf star:bPAC+ brain samples, meanwhile at 6 dpf only 60 of those genes were differentially expressed, and they generally showed a lesser degree of fold change (Supplementary Fig.3d). This suggested that proliferative radial glia might have been lost in the hypothalamus of 13 dpf star:bPAC+ larvae due to cellular senescence. Additionally, we confirmed that the loss of hypothalamic proliferation is maintained into later life, since we observed that in juvenile star:bPAC+ fish, the hypothalamus is smaller and contained fewer mitotic cells (Supplementary Fig.4a-c). This suggests that the proliferative hypothalamic cells lost at 13 dpf do not re-enter the cell cycle later in development, suggesting that they have become senescent, rather than quiescent.

Since we observed a dramatic reduction in cell proliferation in the hypothalamus of 13 dpf star:bPAC+ larvae, we postulated that downstream neurogenesis and neuronal differentiation might also be affected. Since one of the hypothalamus-regulated functions is feeding behaviour and food intake, we sought to determine whether specific neurons that regulate feeding were altered in the hypothalamus of star:bPAC+ larvae at 13 dpf. *agouti-related peptide* (*agrp*) is expressed in a cluster of cells in the arcuate nucleus of the hypothalamus, and we observed significantly fewer numbers of these cells in star:bPAC+ larvae (Fig.3h,i). A small number of cells expressing *cart4* (*cocaine- and amphetamine-regulated transcript*) were observed near to the medial lateral hypothalamus (mLH) and the total number of these cells were also reduced in star:bPAC+ larvae (Fig.3h,j). Orexin+ (*hcrt*, *hypocretin*) cells were observed in the dorso-rostral hypothalamus and we observed a trend reduction in cell count in star:bPAC+ larvae (Fig.3h,k). We observed *npy* (*neuropeptide y*) cells near the mLH and these cells were not affected in star:bPAC+ larvae (Supplementary Fig.5a,b). Finally, *pomca* (*proopiomelanocortin*) is also expressed in the arcuate hypothalamic nucleus, and these cells were not significantly reduced in star:bPAC+ larvae (Supplementary Fig.5a,c). Of the neuromodulators where we observed reduced numbers in star:bPAC+, *agrp*^25, 26^ and *hcrt*^27^ are both known to be appetite-stimulators, meanwhile the effects of *cart* on appetite are more complex^34^. These data indicate that neurogenesis of specific neuronal subtypes that stimulate feeding are reduced in 13 dpf star:bPAC+ larvae.

### Progenitor cells fail to differentiate in the hypothalamus of star:bPAC+ larvae

Since we observed that the effects of GC on hypothalamic proliferation changed dramatically between 5 dpf and 13 dpf, we sought to track the proliferating radial glia of the hypothalamus between these timepoints to determine their fate. To do this we exposed 4 dpf larvae to BrDU (Bromodeoxyuridine) overnight for 17 hours until 5 dpf and then fixed larvae at 5, 8, 10 or 13 dpf for double-IHC analysis of BrDU and Pcna (Fig.4a). At 5 dpf we observed successful incorporation of BrDU into proliferating cells, as observed by co-expression with Pcna, in the hypothalamus of both star:bPAC+ and wild-type larvae (Fig.4b). By 8, 10 and 13 dpf, BrDU+ cells in the wild-type hypothalamus have begun exiting the cell cycle and are now mostly observed around the perimeter of the Pcna-expressing proliferative zone (Fig.4c, Supplementary Fig.6). Indeed, we observed that the percentage of hypothalamic BrDU+ cells that co-express Pcna drops from around 75% at 5 dpf to 50-55% at 8-13 dpf (Fig.4d), suggesting that some of the BrDU+ cells that were proliferating at 5 dpf have lost their progenitor cell status and differentiated.

**Figure 4.**
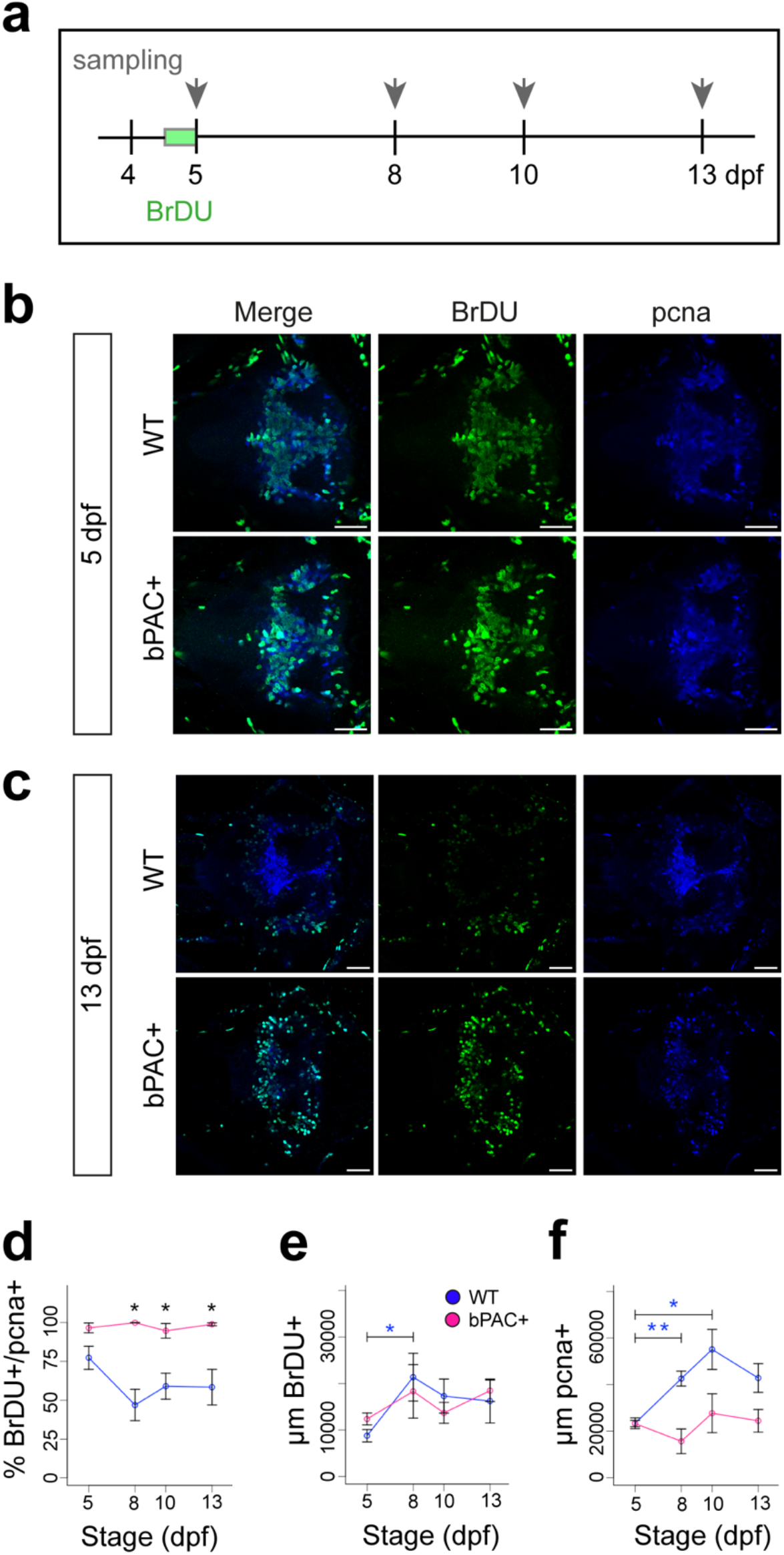
Progenitor cells fail to differentiate in the hypothalamus following elevated GC. **a**. Schematic showing the experimental paradigm. Wild-type and star:bPAC+ larvae were treated with BrDU overnight from 4 dpf until 5 dpf for 17 hours. Following removal of BrDU compound, larvae were collected at 5 dpf, 8 dpf, 10 dpf and 13 dpf for lineage tracing analysis. **b,c.** Confocal images of BrDU lineage tracing analysis at 5 dpf (b) and 13 dpf (c). In each panel wild types are shown in the upper row and star:bPAC+ larvae in the bottom row. The left-most column (Merge) shows the overlap between BrDU-labelled cells with endogenous pcna, whilst single channel images for BrDU and pcna are adjacent. All images are a single plane from a confocal z-stack. In all cases the scale bar indicates 40 μm. **d-f.** Quantitative analysis of BrDU cell fate. Whilst the percentage of BrDU cells expressing pcna decreases over time in wild types, this remains high in star:bPAC+ (d). Whilst BrDU+ cells increase between 5 dpf and 8 dpf in wild types, this does not occur in star:bPAC+ (e). Similarly, pcna+ cells increase between 5 dpf and 10 dpf in wild types, but not in star:bPAC+ larvae (f). Wild types are shown in blue, star:bPAC+ in pink. All graphs show the mean with standard error. N=5-10 per group. *, p<0.05, **, p<0.01 for t-tests with Bonferroni correction.

In stark contrast, hypothalamic BrDU+ cells in star:bPAC+ larvae are still predominantly observed within the Pcna+ proliferative zone after 5 dpf (Fig.4c, Supplementary Fig.6). In fact, the percentage of hypothalamic BrDU+ cells that co-express Pcna remains at around 100% from 5 dpf until 13 dpf in star:bPAC+ larvae (Fig.4d). This indicates that in star:bPAC+ larvae, hypothalamic progenitor cells remain proliferative between 5 dpf and 13 dpf and do not differentiate. Further, we observed an increase in the number of hypothalamic BrDU+ cells between 5 dpf and 8 dpf in wild types, supporting that these cells are dividing and therefore increase in number, however no such increase was observed in star:bPAC+ larvae (Fig.4e). Similarly, the number of hypothalamic Pcna+ cells increased in wild types between 5 dpf and 10 dpf, indicating that the proliferative zone continues to expand as cells divide during this period (Fig.4f). No increase in the number of hypothalamic Pcna+ cells was observed in star:bPAC+ larvae during this developmental period (Fig.4f), further supporting that neurogenesis appears to stall during this developmental window in star:bPAC+ larvae. Together, these data support that in star:bPAC+ larvae, hypothalamic progenitor cells fail to differentiate, and neurogenesis is stalled during the 5-13 dpf time window.

### Functional consequences of elevated GC: precocious feeding followed by physical decline

The increased size and proliferative capacity of the hypothalamus in 5 dpf star:bPAC+ larvae suggested that development of the hypothalamus is accelerated in these animals. Therefore, we hypothesised that behaviours associated with the hypothalamus that develop around this developmental stage might also be altered and thus we analysed feeding in 5 dpf larvae. Feeding usually emerges at around 120 hpf (5 dpf) in lab-raised zebrafish larvae, and larvae are capable of eating small live food. We exposed 5 dpf larvae to live rotifers and saw that star:bPAC+ larvae consumed significantly more rotifers during the trial (Fig.5a), suggesting that GC-induced precocious hypothalamic development is accompanied by early emergence of an associated behaviour, feeding.

**Figure 5.**
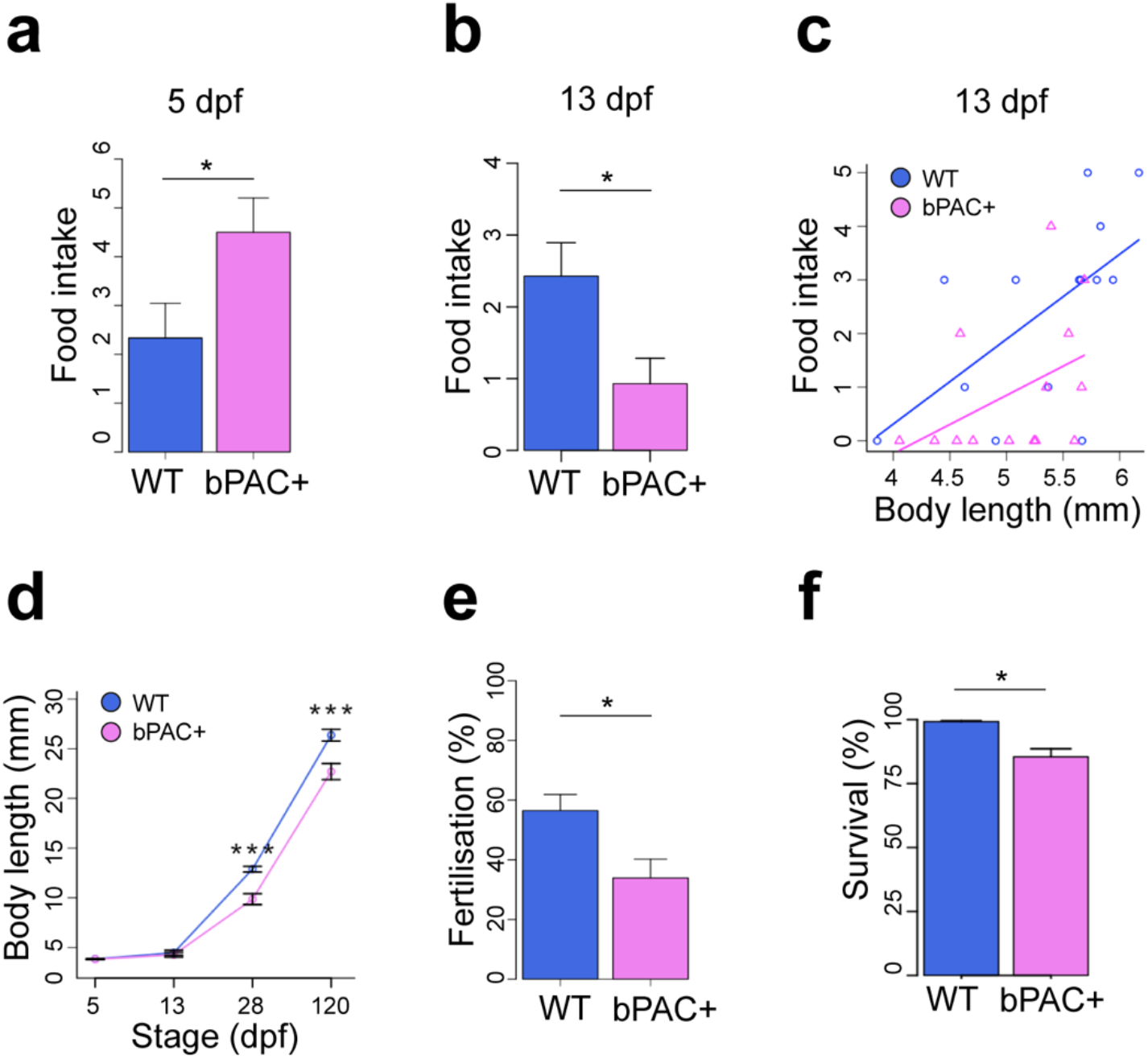
Functional consequences of elevated GC: precocious feeding followed by physical decline. **a.** Analysis of feeding behaviour reveals that 5 dpf star:bPAC+ larvae eat more (N=12, t = -2.1694, df = 21.996, p-value = 0.04113). **b.** Analysis of feeding behaviour in 13 dpf wild-type and star:bPAC+ larvae reveals that food intake is significantly reduced (N=14, t = 2.5626, df = 24.299, p-value = 0.017). **c.** In 13 dpf wild-type larvae, food intake correlates positively with body size (linear regression, R-squared: 0.3619, F: 6.805 on 1 and 12 DF, p-value: 0.02286), however in star:bPAC+, there is no significant correlation (R-squared: 0.1877, F: 2.773 on 1 and 12 DF, p-value: 0.1218). **d.** star:bPAC+ larvae are significantly smaller than wild types at 28 dpf and 120 dpf, but not at 5 dpf or 13 dpf (Two-way ANOVA with Tukey’s test, *** p<0.001 for 28 and 120 dpf). **e**. Percentage of embryos that were fertilised by adult male star:bPAC+ fish is significantly lower compared to wild types (N=16,19 pairs, t = 2.7181, df = 31.072, p-value = 0.01064). **f.** Percentage of star:bPAC+ fish that survived to 2 months is significantly reduced, compared with wild types (Wilcoxon test, N=9, W = 65.5, p-value = 0.004315).

In light of the reduction of cells expressing feeding-related neuromodulators in the hypothalamus of 13 dpf star:bPAC larvae, we sought to determine whether feeding was also affected at 13 dpf. In stark contrast to 5 dpf, at 13 dpf star:bPAC+ larvae fed less than wild types (Fig.5b). Interestingly, in 13 dpf wild-type larvae we observed that food intake positively correlated with body size, in that larger larvae ate more, whilst in star:bPAC+ larvae no significant trend was apparent. (Fig.5c). Hence, star:bPAC+ larvae consumed less, regardless of their size. Whilst at 5 dpf and 13 dpf we did not observe any difference in larval body size, at 28 dpf and 120 dpf star:bPAC+ fish were significantly smaller than wild types (Fig.5d). This suggests that the reduction in food intake in star:bPAC+ animals from 13 dpf ultimately leads to impaired growth. Finally, we also observed reduced fertilisation rates and reduced long-term survival of star:bPAC+ fish (Fig.5e,f). Together, these results point towards potential functional consequences of failed hypothalamic development after 5 dpf, including impaired behaviour and an early physical decline. Further work is required to directly link altered hypothalamic neurogenesis with the observed functional alterations in GC-exposed animals.

## Discussion

This work shows for the first time that elevated glucocorticoid alters the trajectory of hypothalamic development and function. We identify that the hypothalamus is a highly GC-sensitive region where elevated GC causes precocious development followed by failed maturation and early decline accompanied by impaired feeding, growth, and longevity.

Whilst others studies have reported GC-induced reduction in cell proliferation^35^, our data supports the hypothesis that, at least in some cases, the reduced cell proliferation following GC exposure might be a consequence of a prior increase in proliferation. Indeed the concept of GC-inducible stem cells is not novel^36, 37^, and there are many reports of a GC-induced increase in cell proliferation of various cell types including NSPCs^38–41^. The initial GC-induced increase in cell proliferation observed here might be in part mediated by mechanisms akin to those observed during brain injury, whereby formerly quiescent radial glia enter the cell cycle in response to trauma^32, 42^. In the published literature, there are very few previous studies of effects of GC on cell proliferation across development. Of those, Noorlander *et al*., observed that antenatal GC treatment in mice led to an initial reduction in cell proliferation in the embryonic hippocampus, followed by increased cell proliferation during the postnatal period and a subsequent reduction in proliferation during adulthood^43^. There are also examples in mouse models where ELS induced an initial increase in cell proliferation in the hippocampus during early life which was again followed by a subsequent reduction in cell proliferation^10^ or hippocampal volume^12^ in later life. Further, a meta-analysis of age-dependent effects of ELS on cell proliferation shows a negative correlation between age and change in cell proliferation, with the ELS-exposed animals analysed earliest in development exhibiting increased cell proliferation and older ELS-exposed animals exhibiting a reduction in cell proliferation^44^. It is possible that our interpretation of the results presented here also applies to other studies, in that GC drives precocious proliferation leading to failed maturation and a rapid decline. In human hippocampal progenitor cells, a low dose of GC was found to enhance cell proliferation, via MR function, meanwhile a higher dose of GC was reported to diminish cell proliferation, via GR function^45^. Whilst these results support a dichotomy of effects on proliferation resulting from low and high dose GC treatment, it is plausible that a similar phenomenon might occur in response to short-term and long-term exposure to GC, as is seen in our contrasting data from the 5 dpf and 13 dpf GC-exposed hypothalamus. Indeed, further work *in vitro* supports that the effects of GC on neurogenesis can vary depending on the timing of GC treatment^46^.

In our model, we observed profound effects on a population of NSPCs within the proliferative zone of the developing hypothalamus. Indeed this population of hypothalamic radial glia might be akin to tanycyte cells observed in mammals, which also express *Rax*/*rx3* and *Blbp*^18^, and were recently shown to be sensitive to ELS exposure in adult mice^17^. We postulate that these cells might be especially sensitive to the effects of elevated GC, since they are adjacent to the 3^rd^ ventricle and thus might detect elevated GC levels within the CSF. Indeed, neurogenesis of tanycyte cells is even responsive to dietary cues^47^. Another potential driver of increased sensitivity of the hypothalamus to GC would be increased levels of GR, however, it is not currently clear whether the hypothalamus expresses higher levels of GR than other brain regions in zebrafish, or indeed in other vertebrates. As such, an explanation for the specificity of the proliferation phenotype observed in our study remains elusive. We cannot rule-out the possibility that the hypothalamus is not uniquely affected in our model, but rather that the effects of GC on different brain regions might emerge across different time-scales.

We observed significant effects on feeding behaviour of developmental GC-exposed star:bPAC+ larvae, whereby 5 dpf larvae consumed more and 13 dpf larvae consumed less than controls. The observed reduction in feeding persists into adulthood in this model^19^. These changes correlated with increased proliferative hypothalamic radial glia at 5 dpf, which were reduced by 13 dpf, alongside a reduction in numbers of feeding-regulating neurons that express *agrp*, *cart* and *hcrt at* 13 dpf. The role of these hypothalamic neuropeptides in food consumption in mammals is well established^48–50^. In larval zebrafish, hypothalamic *agrp* neurons are known to stimulate feeding^25, 26^, and orexin, produced by *hcrt* neurons, is also known to stimulate food intake in zebrafish^27^. As such, it is likely that the observed reduction in numbers of *agrp* and *hcrt* neurons in 13 dpf star:bPAC+ larvae underlie the observed reduction in food consumption at the same developmental stage. Another hypothesis regarding the basis of the altered food consumption in the star:bPAC+ larvae relates to the observed changes in hypothalamic radial glia. The population of hypothalamic radial glia altered by developmental GC exposure in our model display similarities to mammalian tanycyte cells, based on location, morphology, and marker expression^18^. Recent work demonstrated that hypothalamic tanycytes can regulate food intake^51^. Hypothalamic tanycytes are known to project to neighbouring orexigenic *npy*/*agrp* and anorexigenic *pomc* neurons of the arcuate nucleus, and tanycyte activation was shown to depolarise these neurons *in vitro*, and lead to hyperphagia *in vivo*. It is interesting to postulate that if similar functionality occurs in the hypothalamic radial glia of zebrafish, then the observed increase in these cells at 5 dpf might contribute to the increased feeding behaviour observed, meanwhile the subsequent reduction of these cells might contribute to the reduction of feeding behaviour observed in 13 dpf star:bPAC+ larvae. Further work is required to characterise the development and function of the zebrafish hypothalamic radial glia, and to determine the mechanisms underlying the altered feeding observed in star:bPAC+ larvae.

We hypothesise that the precocious hypothalamic development observed here may represent an example of GC-induced adaptive plasticity (Fig.6), as observed in other animals. For example in red squirrels, females exposed to high-density cues (mimicking increased competition) increase their levels of endogenous GC which drives them to produce offspring that grow faster than controls^52^. This so-called adaptive plasticity is acknowledged to have short-term fitness benefits, but the investment is often associated with costs in later life^53^. The costs of accelerated development can include reduced longevity, which is known as the growth-rate lifespan trade-off^54^, as well as reductions in breeding success^55^. Interestingly, there is evidence directly linking divergence from the normal growth trajectory with longevity, since sticklebacks that are forced to grow faster have a reduced lifespan, whilst those in which growth was slowed down lived longer than controls^54^. In our model, chronic GC exposure ultimately appears to lead to allostatic overload, a maladaptive state whereby an organism can no longer adapt to its conditions^56^. Whilst GC-induced acceleration of growth-rate leading to adverse phenotypes later is widely observed at the organism level, the underlying cellular and molecular mechanisms are poorly understood. Our work provides new insight into these processes in the developing brain.

**Figure 6.**
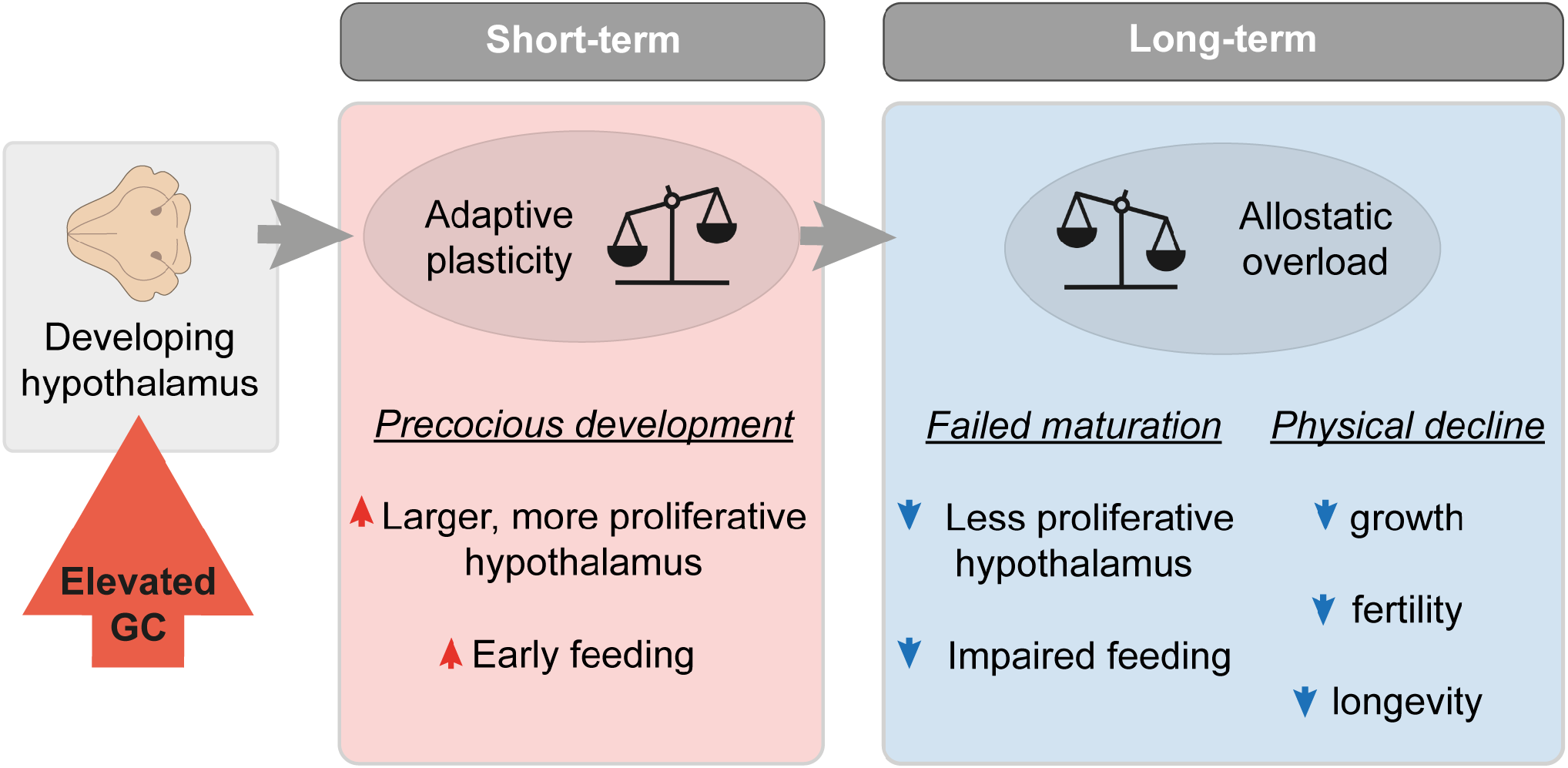
Model proposing how elevated GC alters the trajectory of hypothalamic development and function. In our model of elevated GC, we suggest that in the short-term, GC exposure drives adaptive plasticity, whereby we observed precocious hypothalamic development. In the long-term, chronic GC exposure leads to an accumulation of allostatic load, leading to allostatic overload. We observed this via failed hypothalamic maturation during the larval stage, and subsequent early physical decline of exposed fish.

In the context of the early-late life trade-off, the subsequent reduction in breeding success and/or longevity is thought to be related to oxidative stress^57, 58^. Exposure to elevated GC is known to increase oxidative stress^59^, a phenomenon in which an over-accumulation of reactive oxygen species (ROS) occurs in cells and tissues^60^. ROS is known to cause cellular damage, especially to DNA, and can lead to accelerated telomere shortening^61^. In a previous study of amphibian larvae, the relationship between stress, development, growth and aging depended on the type of stressor exposure^62^. Predator exposure initiated faster development and enhanced growth of survivors, but they showed signs of oxidative stress, had shorter telomeres and reduced long-term survival^62^. This indicates that stress exposure may initiate faster development in early life, but at a cost to long-term health and longevity.

Telomeres shorten with every round of cell division and when a critical size is reached this imposes a functional limit on cell replication, leading to replicative senescence, which can contribute to age-related diseases^61^. One possibility in our model is that the excess proliferation of hypothalamic progenitors observed in 5 dpf star:bPAC+ larvae leads to subsequent senescence of those cells. In contrast to quiescent, apoptotic, or terminally differentiated cells, senescent cells have incurred an irreversible cell cycle arrest, yet remain viable, have alterations in metabolic activity and undergo dramatic changes in gene expression^63^. In support of our hypothesis, a previous study of liver progenitor cells reported that GC exposure induced cell proliferation leading to long-term replicative senescence/stemness exhaustion^64^, and GC is also reported to induce senescence of NSPCs *in vitro*^65^. The presence of senescent cells in the proliferative zone of the GC-exposed hypothalamus at 13 dpf might explain why we observed reduced expression of Pcna and *rx3*, and a reduced proliferative capacity of the remaining glial cells in the proliferative zone of star:bPAC+ larvae. Telomere shortening and senescence are associated with aging and age-related diseases; and models of accelerated aging present with phenotypes such as reduced longevity and physical decline^66^, as observed in our model. Additionally, chronic exposure to GC is associated with signs of accelerated aging at a cellular level^67^. Further analysis of oxidative stress and telomere homeostasis would provide further insight into the mechanisms underlying the phenotypes observed here.

We conclude that the phenotypes observed in our optogenetic model are the result of exposure to elevated GC produced in the interrenal, rather than to other interrenal steroid hormones, which are not detectable in larval zebrafish^19, 68, 69^. We propose that GC-induced alteration of the hypothalamic development trajectory observed in our model is mediated by GR-regulated gene transcription and potentially epigenetic mechanisms. In support of this, we showed that treatment with GR antagonist RU-486 reduced the number of pH3+ mitotic cells in the hypothalamus of star:bPAC+ larvae. We also provide evidence for regulation of hypothalamic *rx3* by GR. Indeed, GR signalling is known to regulate cell cycle progression in NSPCs by inducing a specific pattern of DNA methylation during aging^70^. In a model of replicative senescence, GC exposure lead to altered methylation of GR target gene *fkbp5*, which was exacerbated by age, and the subsequently increased *fkbp5* expression was associated with inflammation and myocardial infarction in humans, suggesting that GC is linked with age-related disease via a mechanism involving epigenetic regulation of *fkbp5*^71^. Meanwhile in cancer cells, GC exposure can induce a cell dormancy state which is in part senescence and in part quiescence, mediated by GR target gene *CDNK1C* (*cyclin-dependent kinase inhibitor 1C)*^72^, further supporting that GC exposure might regulate cell proliferation in our model via GR-mediated regulation of cell cycle.

We speculate that the effects of elevated GC on the hypothalamic development trajectory reported here might be an example of allostatic overload. Allostasis is the process of activating adaptive mechanisms in response to external or internal changes, however prolonged exposure to stressors can lead to accumulation of so-called allostatic load, and subsequently a maladaptive state of allostatic overload^8^, which is associated with age-related diseases and reduced longevity. In support of this, it has been proposed that effects of stress or GC on NSPCs in young individuals may affect their renewal potential in the long-term, predisposing to adult disease^36^. We hypothesise that whilst precocious hypothalamic development induced by GC might be an example of adaptive plasticity in the short-term, the cost of this may be paid in later life through subsequent replication senescence of those cells, leading to failed hypothalamic maturation, subsequent impaired feeding behaviour and ultimately physical decline (Fig.6). Further understanding of how stress and GC exposure can alter development trajectories at the molecular and cellular level is of critical importance to reduce the burden of mental and physical ill health across the life-course.

## Methods

### Zebrafish Husbandry and Maintenance

Zebrafish experiments were performed at the University of Exeter Aquatic Resource Centre and at the Johannes Gutenberg University of Mainz in compliance with local and national animal welfare laws, guidelines, and policies and approved by local government (Landesuntersuchugsamt Rheinland-Pfalz, Germany – 23 177-07/G20-1-033 and Home Office PPL number: PEF291C4D). *Tg(2kbStAR:bPAC-2A-tdTomato)^uex^*^300^ were bred with wild-type TU zebrafish strain and were screened at 4 dpf on a fluorescent stereomicroscope. Adult zebrafish were maintained under standard zebrafish husbandry conditions, under a 12:12 light/dark cycle^73^.

### RNA sequencing

RNA sequencing data was obtained from our other study^19^, which is deposited in the European Nucleotide Archive (ENA, PRJEB53713). The detailed protocol for sample collection, RNA preparation, mRNA sequencing, and bioinformatic analysis is available in an online methods repository^74^.

### Gene Ontology analysis

The functional enrichment analysis was performed using g:Profiler (version *e107_eg54_p17_bf42210)* with an adjusted p-value of 0.05. For GO analysis of neurogenesis gene clusters (Fig1e), significant GO terms for each cluster were manually slimmed according to QuickGO ancestor charts. Heatmaps were generated in R using the heatmap.2 function.

### qPCR

Fish were immobilised using ice-cold water and larvae or dissected juvenile/adult brains were stored in RNA later solution. Sample collection was carried out between 08:00 and 10:00. Larvae were subsequently dissected, such that each replicate consisted of 15 larval heads (eyes and jaw removed) or 3 juvenile/adult brains. RNA was extracted using TRIzol™ Reagent (Invitrogen™, 15596026), as previously described^75^, and cDNA was synthesised using High-Capacity RNA-to-cDNA™ Kit (Applied Biosystems, 4387406). Approximately 100 ng cDNA was used in a 10 μl qPCR reaction with PowerUp™ SYBR™ Green Master Mix (A25778, Applied Biosystems). Primer sequences can be found in Supplementary table 1. The reactions were run in Hard-Shell 96-Well PCR Plates (HSP9601, BioRad) on a CFX96 Real Time PCR machine (Bio-Rad) using a standard protocol. Relative expression was calculated using the 2^−ΔΔCt^ method, with *18s* as a reference gene.

### Fluorescent in situ hybridisation (FISH) and immunohistochemistry (IHC)

The FISH and IHC methods were based on a protocol by Jakob von Trotha (2016) for whole zebrafish embryos and larvae. Larvae or juvenile fish were immobilised in ice-cold water and then whole larvae or dissected juvenile fish brains were fixed in 4% PFA overnight at 4 °C, dehydrated in methanol and subsequently stored at -20C. After rehydration, larvae or brains were permeabilised using proteinase K (10 μg/ml) treatment at 37 °C and then re-fixed in PFA. DIG-conjugated mRNA probes were hybridised overnight at 65 °C. *In situ* hybridisation probes for *rx3*, *agrp*, *cart4, npy* were synthesised using primers listed in Supplementary Table 1. *pomc* probe was synthesised from a plasmid, as previously described^76^. After removal of probe and washing, samples were blocked for 2 hours in blocking solution (1% blocking reagent; (Roche, 11096176001) in malic acid buffer (0.15M maleic acid, 0.15M NaCl, PH 7.5)) and then incubated overnight at 4 °C in anti-DIG POD antibody (Roche, 11207733910) diluted 1:300 in blocking solution. After more washing, samples were incubated for 40 minutes in the dark with 1:200 FITC-tyramide (synthesised from product 46410, Thermo Scientific), 0.003% H2O2, 2% dextran sulphate in PBST. For IHC antigen retrieval was performed according to published methods^77^. Samples were blocked in 10% Normal goat serum in PBST for 2 hours before incubation in primary antibody overnight at 4 °C. For IHC of BrDU-treated samples, samples were treated with 1N HCl for 30 minutes prior to blocking. Primary antibodies used were: pH3 (Merck/Millipore, 06-570), pcna (Sigma, MABE288), BLBP (Sigma, ABN14), otpa^78^, BrDU (abcam, ab6326). After washing all day, samples were incubated overnight in 1:1000 secondary antibody with 1:200 hoechst (H3570, Invitrogen). Secondary antibodies used were Goat anti-Rabbit IgG (H+L) Cross-Adsorbed Secondary Antibody, Alexa Fluor™ 488 (A-11008, Invitrogen), Goat anti-Rabbit IgG (H+L) Cross-Adsorbed Secondary Antibody, Alexa Fluor 633 (A-21070, Invitrogen), Goat anti-Mouse IgG (H+L) Cross-Adsorbed Secondary Antibody, Alexa Fluor 568 (A-11004, Invitrogen), Goat anti-Rat IgG (H+L) Cross-Adsorbed Secondary Antibody, Alexa Fluor 488 (A-11006, Invitrogen). After staining all samples were washed further then cleared overnight in glycerol. Samples were imaged on a Zeiss LSM 880 confocal microscope using a 25x or 10x objective.

### Image processing

Image processing was performed in Fiji (ImageJ2, version 2.9.0). For whole brain counting of cells, z-stacks were taken across the brain from dorsal to ventral and regions were defined using the hoechst staining with the Z-Brain viewer^79^. Images were thresholded and PH3 cells in each region were manually counted in a blinded manner. For counting of cells within the hypothalamus, cell counts were performed manually from z-stacks, in a blinded manner. Similarly, hypothalamic volume was calculated based on hoechst staining in z-stacks, using Z-Brain viewer. Corrected Total Cell Fluorescence (CTCF) was calculated as Integrated Density of cells – (Area of cells X Mean fluorescence of background reading). For CTCF calculations, labelled cells were identified first by thresholding each image and creating ROIs of the labelled cells using the Analyze Particles function. The total Integrated density and total area of all labelled cells was then used to calculate CTCF for each sample. For co-expression analysis, ROIs of labelled cells were generated for each marker as described above. Percentage overlap was calculated using the Analyze Particles function with ROIs of marker 1 cells against a mask of marker 2 cells. Total area of cells was calculated from the thresholded mask/ROIs of labelled cells. pH3 cells were manually scored as *rx3*- or *rx3*+ using the ROIs/masks as described above, in a blinded manner. All measures are normalised relative to size of the hypothalamus.

### Feeding behaviour

Prior to behavioural analysis, larvae were transferred to 24 well cell culture plates (83.3922, Sarstedt) with 1 larva per well in 1.5 ml aquarium water. Larvae were then left to acclimate to the behavioural testing room (maintained at 28 °C) for 2 hours. 5 dpf larvae were fed 1 ml of diluted live rotifers (approximately 20), meanwhile 13 dpf larvae were fed 5 live artemia. Larvae were recorded for 10 minutes using Basler Video Recording Software. The camera used was a Basler (Germany) acA1300-200um USB 3.0 camera with ON Semiconductor PYTHON 1300 CMOS sensor (203 frames per second at 1.3 MP resolution) with a Computar Zoom Lens 18-108/2.5 (Japan). Number of successful and unsuccessful prey captures and latency to first hunting attempt were analysed manually from video recordings in a blinded manner.

### BrDU experiment

Larvae were transferred to a 12 well dish with 10 larvae per well. Larvae were incubated overnight from 4 dpf in 3 ml 10 mM 5-Bromo-2′-deoxyuridine (B5002, Sigma) with 1% DMSO for 17 hours. On the morning of 5 dpf, BrDU was removed by washing 3-times with aquarium water. Larvae were subsequently fixed in PFA or raised until a later stage for fixation.

### Drug treatments

Larvae were incubated from 6 hpf until 120 hpf with 50 μM dexamethasone (Sigma-Aldrich D2915) or from 48 hpf until 120 hpf with 2 μM RU-486 (Sigma-Aldrich, M8046). DMSO solution was used as a control. Solutions were changed daily.

### ChIP-qPCR

We analysed the sequence for the *rx3* gene (ENSDARG00000052893, chromosome:GRCz11:21:10755554:10759823:1). Primers for *rx3* GRE1 and GRE2 are described in supplementary table 1 and were designed using Primer3. GR bounded chromatin was prepared following the protocol from Idilli *et al*., 2020^80^, with minor modifications. For each sample, 100 dissected 5 dpf larval heads (without eyes) were incubated for 10 minutes at room temperature in 1% formaldehyde in PBS with protease inhibitors (A32955, Thermo Scientific) for cross-linking. We used a glucocorticoid receptor polyclonal antibody (24050-1-AP, Proteintech) and Rabbit IgG Isotype Control (10500C, Invitrogen) as primary antibodies.

### TUNEL assay

Apoptotic cells were labelled using the In Situ Cell Death Detection Kit, Fluorescein (11684795910, Roche). Larvae were first fixed, dehydrated, rehydrated and permeabilised as per the FISH protocol. Larvae were incubated in TUNEL reaction mix at 37 °C for 2 hours and subsequently, washed and cleared as per the FISH protocol.

### Statistics

All analyses and graphs were generated in R (RStudio 2022.07.2). Prior to testing for statistically significant differences between groups, data were tested for normality and variance. Where data did not fit the assumptions for t-testing or ANOVA, non-parametric alternatives were used. Where appropriate, Bonferroni correction for multiple testing was used. In all bar graphs mean +/- standard error is presented for each group.

## Data Availability

All sequenced reads for RNA-seq were deposited in European Nucleotide Archive (ENA, PRJEB53713).

## Author contributions

HE designed the study, performed the experiments, analysed the data, provided funding and wrote the manuscript. MK designed, performed and analysed the RNA-seq experiments, and performed the GR-ChIP. AT, JK, and MH performed some of the experiments. SR provided funding and contributed to study design and manuscript writing.

## Acknowledgements

This project is supported by an award from The Dennis and Mireille Gillings Foundation to SR and the German Federal Office for Education Research (grant number 01GQ1404) to SR. HE received support from the Society for Endocrinology and Wellcome Trust Institutional Strategic Support Fund 3 scheme (ISSF3) to Translational Research Exchange @ Exeter. MH received support from the CN Yang Scholars Programme at Nanyang Technological University. We would like to acknowledge the support of Ms. Kathrin Domdera at the University of Mainz and the Exeter Aquatic Resources Centre staff at the University of Exeter for expert zebrafish care as well as Dr Corin Liddle at Exeter Bioimaging Centre for microscopy. We are grateful to Dr Elina Jacobs, Dr Kate Ellacott, Dr Steffen Scholpp, and Professor Gil Levkowitz for feedback on the manuscript.

