## Supplemental Figures for "Elevated glucocorticoid alters the developmental dynamics of hypothalamic neurogenesis"

**Supplementary Figure 1**

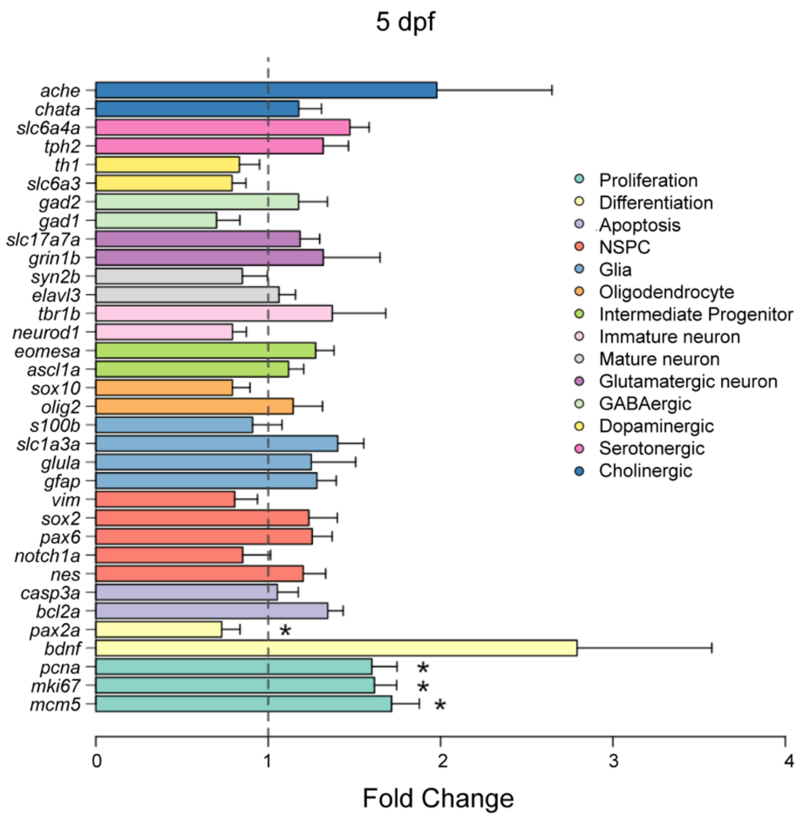


**Cell proliferation is increased in brains of 5 dpf star:bPAC+ larvae.**

qPCR analysis of 5 dpf whole brain samples indicates a significant increase in mRNA expression of three genes associated with cell proliferation (*pcna*, *mki67*, *mcm5*) and one gene associated with cell differentiation (*pax2a*) in star:bPAC+ whole brain samples, compared with wild types. Fold change expression (relative to reference gene *sep15*) in star:bPAC+ relative to wild types is shown (N=8 pools of 12 heads/sample, * p<0.05).

**Supplementary Figure 2**

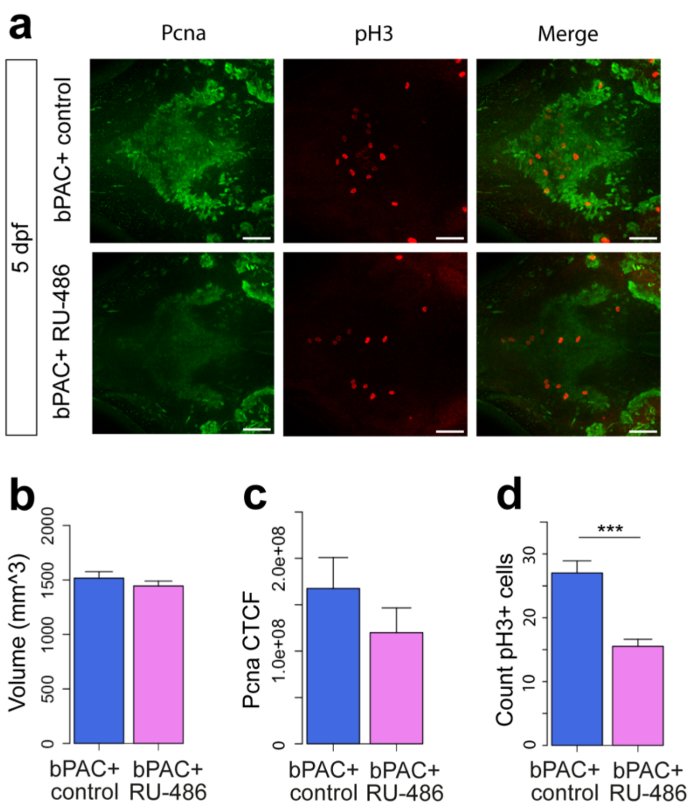


**Hypothalamic proliferation is regulated by GC during development.**

**a.** Maximum intensity projection of a confocal stack through the hypothalamus of 5 dpf star:bPAC+ control fish and following treatment with RU-486, with pH3 (red) and Pcna IHC (green). Scale bar is 40 $\mu$m. **b.** Hypothalamus volume is not affected by RU-486 treatment in 5 dpf star:bPAC+ larvae (t-test, t = 0.96213, df = 16.73, p-value = 0.3497). **c.** Normalised Pcna CTCF in the hypothalamus is not significantly affected by RU-486 treatment in star:bPAC+ larvae (t-test, t = 1.1162, df = 17.165, p-value = 0.2797). **d.** Normalised count of pH3+ cells is significantly reduced by RU-486 treatment in the hypothalamus of 5 dpf star:bPAC+ fish (t-test, t = 5.2469, df = 14.415, p-value = 0.0001122). Graphs show the mean with standard error. N=10. ***, p<0.001.

**Supplementary Figure 3**

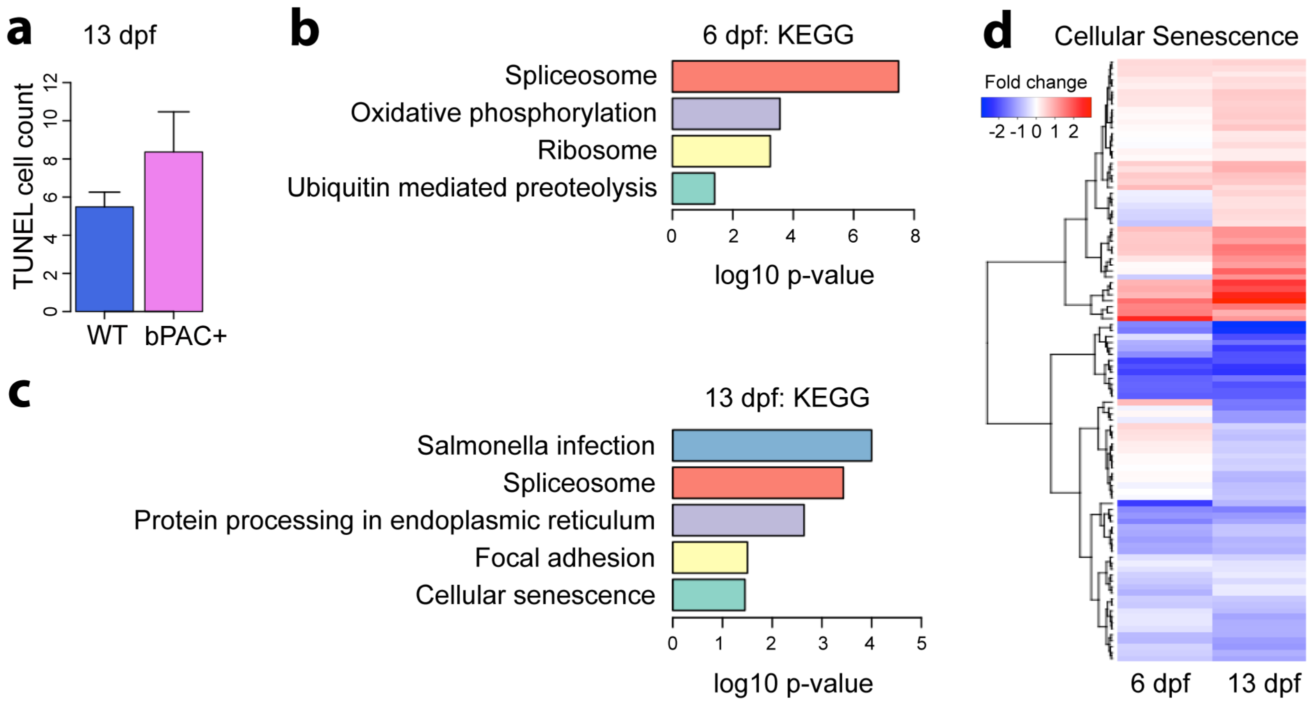


**Cell death and cellular senescence in star:bPAC+ larvae.**

**a.** The total number of TUNEL+ cells in the hypothalamus is not different in 13 dpf star:bPAC+ larvae, compared with wild types (Wilcoxon test, N=6-9, W = 16, p-value = 0.2238). **b-c**. KEGG pathway analysis of DEGs in the RNA-seq data from 6 dpf (b) and 13 dpf (c) star:bPAC+ versus wild-type larvae. Significant pathways with an adjusted p-value of <0.05 are shown. Cellular senescence genes are enriched in 13 dpf DEGs, but not 6 dpf. **d.** Heatmap of log2 fold changes of 102 DEGs associated with KEGG pathway ‘cellular senescence’ at 13 dpf, and their corresponding expression change at 6 dpf in star:bPAC+ brains compared with wild types.

**Supplemental Figure 4**

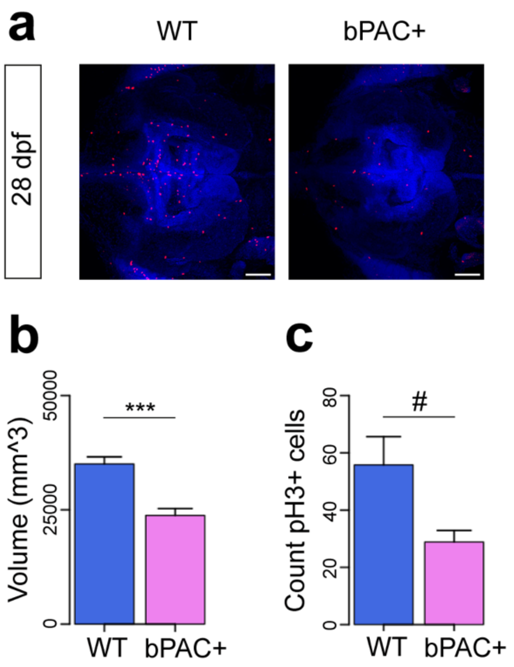


**Hypothalamic proliferation is reduced in juvenile star:bPAC+ fish.**

**a.** Maximum intensity projection of a confocal stack through the hypothalamus of 28 dpf wild-type and star:bPAC+ fish with PH3 IHC (red) and hoechst staining (blue). Scale bar is 100 μm. **b.** Hypothalamus volume is reduced in 28 dpf star:bPAC+ fish (t-test, t = 5.1649, df = 7.993, p-value = 0.0008608). **c.** Normalised count of pH3+ cells showed a trend towards a reduction in the hypothalamus of 28 dpf star:bPAC+ fish (t-test, t = 2.5237, df = 5.3203, p-value = 0.05006). Graphs show the mean with standard error. N=5. #, p<0.1, ***, p<0.001.

**Supplemental Figure 5**

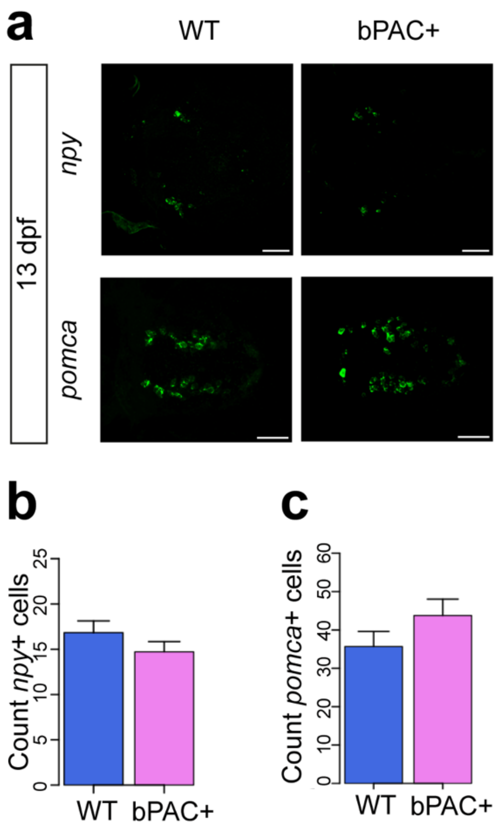


**Expression of feeding-related neuromodulators in the hypothalamus of 13 dpf star:bPAC+ larvae.**

**a.** Expression of genes involved in feeding regulation in the hypothalamus of 13 dpf star:bPAC+ and wild-type larvae as shown in maximum intensity projections of confocal stacks. Scale bar is 40 μm. **b.** The Number of *npy*+ cells is not different (N=18-21, t = 1.231, df = 35.34, p-value = 0.2264 ). **c.** The number of *pomca*+ cells is not significantly different (N=12-16, t = -1.3905, df = 24.592, p-value = 0.1768) in the hypothalamus of 13 dpf star:bPAC+ larvae compared with wild types.

**Supplementary Figure 6**

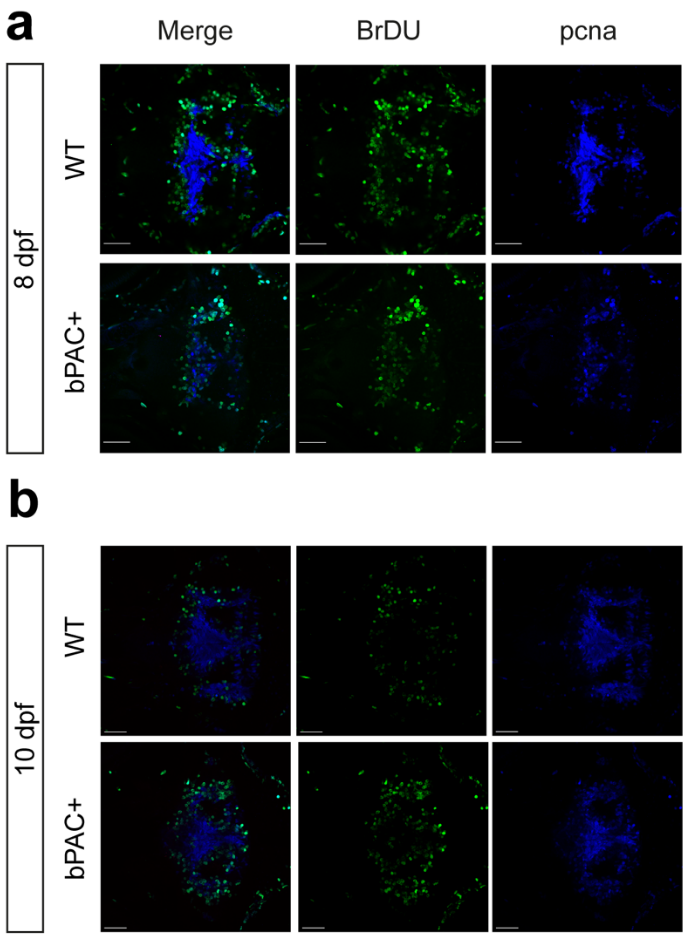


**Altered differentiation in the developing star:bPAC+ hypothalamus.** Confocal images of BrDU lineage tracing analysis at 8 dpf (a) and 10 dpf (b). In each panel wild types are shown in the upper row and star:bPAC+ larvae in the bottom row. The left-most column shows the overlap between BrDU-labelled cells with endogenous pcna, whilst single channel images for BrDU and pcna are adjacent. All images are a single plane from a confocal z-stack. In all cases the scale bar indicates 40 μm.

**Supplementary Table 1**. Primer sequences for qPCR, *in situ* hybridisation probes and ChIP-qPCR.

| Experiment | Name | Sequence | Reference |
|---|---|---|---|
| qPCR | 18s_qF | TCG CTA GTT GGC ATC GTT TAT G | This paper |
| qPCR | 18s_qR | CGG AGG TTC GAA GAC GAT CA | This paper |
| qPCR | ache_qF | GCAAAAGCA TGTGGGCTTGA | This paper |
| qPCR | ache_qR | TCCACTTTCCACTGTCGCTC | This paper |
| qPCR | chata_qF | TGGGGCTGCGAGAGATTGCC | Podlasz 2012 |
| qPCR | chata_qR | TCCTCAGTGGTAGGAACCTGGCT | Podlasz 2012 |
| qPCR | slc6a4a_qF | ACATCTCCTCAAAGCCCCAAA | Jeong 2019 |
| qPCR | slc6a4a_qR | CCACCAGAGTCCTAAATGTTCCA | Jeong 2019 |
| qPCR | tph2_qF | GCAAATACTGGGCTCGGAGA | Sahu et al 2019 |
| qPCR | tph2_qR | GAGCATGGAGGATGCAAGGT | Sahu et al 2019 |
| qPCR | th1_qF | GACGGAAGATGATCGGAGACA | Sahu 2019 |
| qPCR | th1_qR | CCGCCATGTTCCGATTTCT | Sahu 2019 |
| qPCR | slc6a3_qF | CATTTACCCAGAAGCCATTGC | Moriya 2019 |
| qPCR | slc6a3_qR | GCGTGTCGATTCCTAAAGTCA | Moriya 2019 |
| qPCR | gad2-qF | AAGCAGAAGGGATACGTGCC | Cocco 2017 |
| qPCR | gad2-qR | CCGGTGTTTCCGGGACATTA | Cocco 2017 |
| qPCR | gad1_qF | AACTCAGGCGATTGTTGCAT | Hortopan 2010 |
| qPCR | gad1_qR | TGAGGACATTTCCAGCCTTC | Hortopan 2010 |
| qPCR | slc17a7a-qF | GCCAGTGTATGCCATCATCGTAGCC | Liu 2020 |
| qPCR | slc17a7a-qR | TCCACAGTTCATGAGTTTACGGACGTT | Liu 2020 |
| qPCR | grin1b_qF | TTGACAACAAGCGAGGACCC | Gao 2018 |
| qPCR | grin1b_qR | CGTCTTCACTTGCAGACAGGA | Gao 2018 |
| qPCR | elavl3-qF | ATCAACACGCTCAACGGTCT | Ashikawa 2019 |
| qPCR | elavl3-qR | TTACCAGGATGCGTGAGGTG | Ashikawa 2019 |
| qPCR | tbr1b_qpF | CAAGCAGGGACGACGAATGT | This paper |
| qPCR | tbr1b_qR | CATTCGTATCCGCTTTGCCG | This paper |
| qPCR | neurod1_pF1_SP6 | ATT TAG GTG ACA CTA TAGGACAAGTGTCACAGCAGC | This paper |
| qPCR | neurod1_qR1 | GCATCATCGTCGTCTTC | This paper |
| qPCR | eomesa-qF | TAACTAAACAGGGCAGGCGAA | This paper |
| qPCR | eomesa-qR | ATGCGCAGTCAGGTTCAGTC | This paper |
| qPCR | ascl1a_qF | ATCTCCCAAAACTACTCTAATGACATGAACTCTAT | Kaur 2018 |
| qPCR | ascl1a_qR | CAAGCGAGTGCTGATATTTTTAAGTTTCCTTTTAC | Kaur 2018 |
| qPCR | sox10_qF | TCACGCTACAGGTCAGAGTCA | Roberto 2018 |
| qPCR | sox10_qR | ATTTCGCCAATGTCCACG | Roberto 2018 |
| qPCR | olig2_qF | CTTATGCTGAGCAACTCG | This paper |
| qPCR | olig2_qR | AGAGTGCCACAACTGGAC | This paper |
| qPCR | s100b_qF2 | TGCGGATCCAGGTGTTTTAC | Zhang 2019 |
| qPCR | s100b_qR2 | CGCTTTGGTGGGTCTTTTTA | Zhang 2019 |
| qPCR | slc1a3a _qF | GATAGTGGAGATGGACGATCTG | qDB_best |
| qPCR | slc1a3a _qR | TGTCACAAGGAAGTAGAGCAAT | qDB_best |
| qPCR | glula_qF | CAACCTGGAACTCCCACTGAG | Breuer 2019 |
| qPCR | glula_qR | TACTGACGGACACCCCTTTG | Breuer 2019 |
| qPCR | gfap-qF | ACCCGTGACGGAGAGATCAT | This paper |

| | | | |
|---|---|---|---|
| qPCR | gfap-qR | GCCAGTGTCTGAGCCTCATT | This paper |
| qPCR | vim_qF | GGT TCG CCA GTT ACA TAG | This paper |
| qPCR | vim_qR | CTC GTA CAG ATC TCC GAC | This paper |
| qPCR | sox2_qF | CTTCATGGTGTGGTCGAG | This paper |
| qPCR | sox2_qR | CTTCGTCGATGAATGGTC | This paper |
| qPCR | pax6a/b_qF | CAGTACAAGAGGGAGTGTCC | Kleinjan et al., 2008 |
| qPCR | pax6a_qR | CTCACCGCCTCCGTCTGACTGT | Kleinjan et al., 2008 |
| qPCR | notch1a-qF | GGAATATGCGAGTACAAGCCC | Li 2015 |
| qPCR | notch1a-qR | AACACACAGTCGCACTTCAC | Li 2015 |
| qPCR | nes_qF2 | TGGACTGGAGGTGGCAACATACA | Luo 2016 |
| qPCR | nes_qR2 | AGGCAGGAAGCAGCAGTGGTT | Luo 2016 |
| qPCR | casp3a-qF | GAACCGCTTTGTCATCGGAA | This paper |
| qPCR | casp3a-qR | CACCCTTCAACAGCGAGAAC | This paper |
| qPCR | bcl2a_qF | GAGGTTGGGATGCCTTCGTG | Toms 2019 |
| qPCR | bcl2a_qR | CCAAGCCGAGCACTTTTGTT | Toms 2019 |
| qPCR | pax2a_pR1_T3 | AATTAACCCTCACTAAAGGGATCAGTGCTGGTAGCGA G | This paper |
| qPCR | pax2a_qF2 | CTCAACAGCAGCTGGAG | This paper |
| qPCR | bdnf_qF2 | AGAGCGGACGAATATCGCAG | Walter 2019 |
| qPCR | bdnf_qR2 | GTTGGAACTTTACTGTCCAGTCG | Walter 2019 |
| qPCR | pcna_qF | GTCGACAAGGAGGATGAA | This paper |
| qPCR | pcna_qR | GTCTTGGACAGAGGAGTG | This paper |
| qPCR | mki67_qF | TCTCGCATCAGTACCCCATC | This paper |
| qPCR | mki67_qR | ACGGTATCTCTGGTGTTGCA | This paper |
| qPCR | mcm5_qF2 | GCCAGTAGGAGAAGAGACT | Zhang 2017 |
| qPCR | mcm5_qR2 | AGCAGTAGAGGAGATGATGA | Zhang 2017 |
| In situ hybridisation | rx3_sp6_pF2 | ATT TAG GTG ACA CTA TAG GATCCACAGCATTGAGTC | This paper |
| In situ hybridisation | rx3_T7_pR2 | TAA TAC GAC TCA CTA TAG GAACCACACCTGTACTCG | This paper |
| In situ hybridisation | agrp_pF | AGTCTGAGTGATTATGATGCTGA | This paper |
| In situ hybridisation | agrp_T3_pR | AATTAACCCTCACTAAAGGGATCTGTGGATTCTCTGT GCGA | This paper |
| In situ hybridisation | cart4_pF2 | CATTCAGCACCATGGAGAGC | This paper |
| In situ hybridisation | cart4_pR2_T3 | AATTAACCCTCACTAAAGGGAGGTGTCAGTCTCAAGC GTTG | This paper |
| In situ hybridisation | hcrt_pF | AATGCGCCTCTGTCGACATTG | Blechman 2007 |
| In situ hybridisation | hcrt_pR_T3 | AATTAACCCTCACTAAAGGGAACTGCTCACATCCTGT GGTACCG | Blechman 2007 |
| In situ hybridisation | npy_sp6_F | ATTTAGGTGACACTATAGGAAGATGTGGATGAGCTG | This paper |
| In situ hybridisation | npy_T7_R | TAATACGACTCACTATAGTCCTCATATCTGGTCTGG | This paper |
| ChIP-qPCR | rx3_GRE1_qF | CACTGTTGGAGAAAGTCGCT | This paper |
| ChIP-qPCR | rx3_GRE1_qR | ACTGAGATCCAACAAGCCTC | This paper |
| ChIP-qPCR | rx3_GRE2_qF | TGAACATGTTGCACTTTCTCAC | This paper |
| ChIP-qPCR | rx3_GRE2_qR | TCGACCTACAAACTCCTGCA | This paper |
